# Fragile X syndrome-linked mutations impair FMRP accumulation in neuronal transport granules

**DOI:** 10.64898/2026.09.08.750051

**Authors:** Momo Kubo, Maho Ishiyama, Kanae Yoshizawa, Yukio Sasaki

**Author notes:** Corresponding author. E-mail address (Y. Sasaki).

## Abstract

RNA granules are biomolecular condensates that are reversibly formed through multivalent interactions among mRNAs and RNA-binding proteins (RBPs). RNA granules include stress granules and neuronal transport granules, which share common components and functions in translational regulation. Fragile X messenger ribonucleoprotein (FMRP) is an RBP encoded by *FMR1*, whose loss of function causes fragile X syndrome (FXS), a neurodevelopmental disorder. FMRP is a component of both stress granules and neuronal transport granules. Although several FXS-associated missense mutations in FMRP have been identified, it remains unclear whether these mutations affect the assembly of FMRP into RNA granules. Here, we found that the KH1 domain mutation G266E markedly reduced the accumulation of FMRP in stress granules in HEK293T cells, whereas the KH2 domain mutation I304N had little effect. In addition, both G266E and I304N reduced the accumulation of FMRP within neuronal transport granules in FMRP-deficient mouse cortical neurons. Because G266E and I304N have been reported to impair the RNA-binding and/or RNA-associated functions of FMRP, our findings suggest that the KH1 and KH2 domains are important for the assembly of FMRP into RNA granules. These findings further raise the possibility that impaired RNA granule assembly may contribute to the pathogenesis of FXS and other neurodevelopmental disorders.

## Introduction

Neurons are highly compartmentalized, and it is known that specific proteins and mRNAs accumulate into specific compartments, such as neurites and synapses, through liquid-liquid phase separation (LLPS) (Wu et al., 2020; Hayashi et al., 2021; Naskar et al., 2023). LLPS is a phenomenon in which biomolecules aggregate into droplet-like structures, contributing to the formation of structures such as RNA granules and postsynaptic density as “membrane-less organelles”. In particular, RNA granules, composed of RNA and RNA-binding proteins (RBPs), are dynamic ribonucleoprotein assemblies, many of which exhibit properties associated with LLPS. RNA granules facilitate the efficient metabolism, transport, and translational control of mRNAs; well-known examples include stress granules and neuronal transport granules (Hayashi et al., 2021; Naskar et al., 2023; Kiebler and Bauer, 2024). In recent years, mutations in various RBPs have been identified in patients with developmental disorders, and these mutations were found to reduce the formation of stress granules or their accumulation within them (Jia et al., 2022; Bi et al., 2025). However, it remains unclear how aberrant formation of RNA granules is involved in abnormal neuronal development and synaptic pathologies.

Fragile X messenger ribonucleoprotein (FMRP) is an RBP encoded by *FMR1*, whose loss of function causes fragile X syndrome (FXS), a neurodevelopmental disorder (Penagarikano et al., 2007; Bhakar et al., 2012). FMRP is a component of both stress granules and neuronal transport granules (Antar et al., 2004; Lai et al., 2020; Starke et al., 2022). The formation and dissolution of FMRP granules—driven by interactions with mRNAs and other RBPs— constitute a reversible process mediated by LLPS that is regulated by post-translational modification of FMRP (Kim et al., 2019; Prieto et al., 2019; Tsang et al., 2019). Multivalent interactions between FMRP molecules, or between FMRP and other proteins and/or mRNAs, are thought to be crucial for the formation of RNA granules, such as stress granules and neuronal transport granules (Prieto et al., 2019). These multivalent interactions are considered to be mediated by the various domains and regions of FMRP. FMRP consists of two Age domains (Age1, Age2), three KH domains (KH0, KH1, KH2), and an intrinsically disordered region (IDR), which includes the arginine (R)-glycine (G)-rich RGG-box (Fig. 1A) (Prieto et al., 2019). The domains and regions within FMRP are thought to mediate the formation of specific binding interactions and the assembly of RNA granules via LLPS. Specifically, the two Age domains (Age1 and Age2) and the KH0 domain are involved in dimer formation (Adinolfi et al., 2003; Myrick et al., 2014). The KH1 and KH2 domains are involved in RNA binding and are also associated with polysomes (Feng et al., 1997; Ascano et al., 2012; Myrick et al., 2014). The RGG-box interacts with mRNA that forms G-quartet structures, which are higher-order structures of nucleic acids (Blackwell et al., 2010). However, it remains unclear which domains or regions are involved in FMRP-containing RNA granule formation.

**Figure 1.**
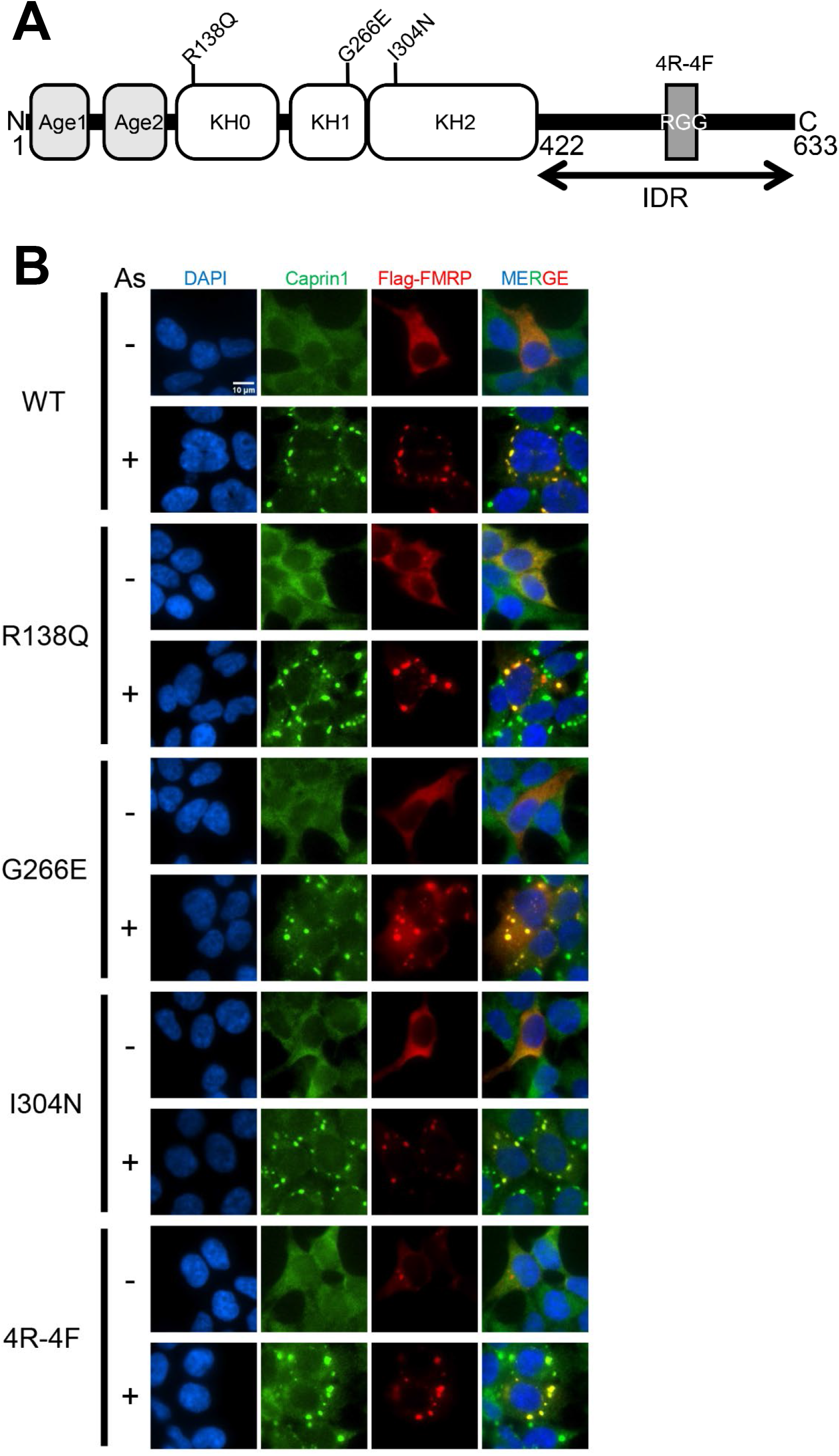
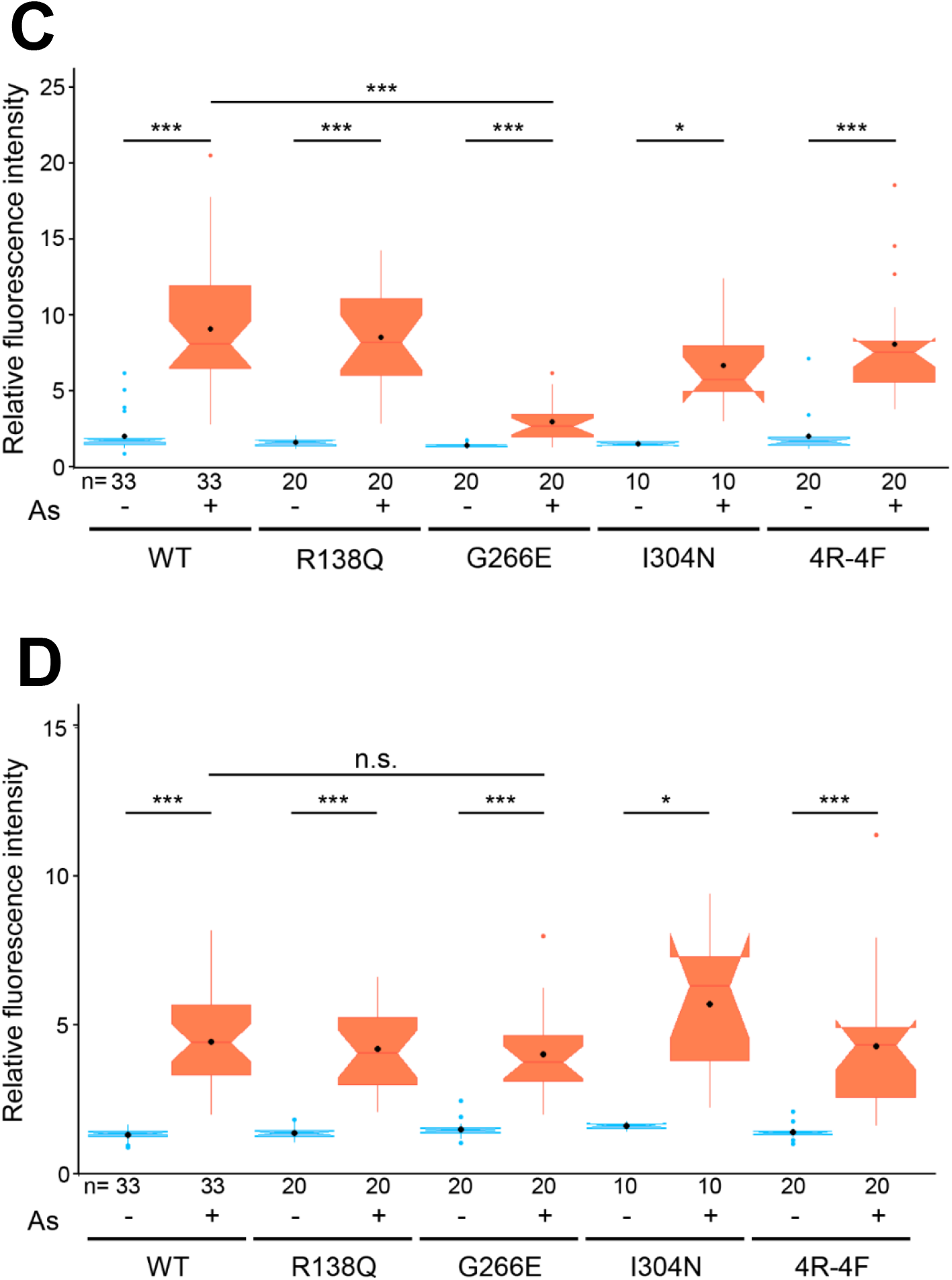
The KH1 domain mutation (G266E) reduced accumulation of FMRP in stress granules. (A) Structure of FMRP domains. FMRP is composed of 2 Age domains, 3 KH domains, and a single RGG box localized at the IDR. Age1, Age2, and KH0 domains are involved in dimer formation. KH1 and KH2 domains are responsible for mRNA binding. RGG box interacts with a G-quadruplex. Patient-derived mutations, R138Q, G266E, and I304N, are indicated. 4R-4F is a mutant in which 4 arginine residues within the RGG box are replaced by 4 phenylalanine residues. (B) Stress granule formation in HEK293T cells expressing FMRP WT or one of the mutants. HEK293T cells were transfected with the plasmid expressing FMRP WT or one of the FMRP mutants (R138Q, G266E, I304N, and 4R-4F). The transfected cells were subjected to oxidative stress treatment (200 μM As) for 1 h, followed by fluorescent immunostaining using anti-Flag and Caprin1 (stress granule marker) antibodies. Scale bar: 10 μm. (C) Analysis of fluorescent intensities of FMRP-containing granules. Fluorescent intensities of FMRP-containing granules, the cytoplasm (excluding the nucleus and granules), and background in (B) were measured. Granules were defined using the “Analyze Particles” tool on binarized images by ImageJ (see Materials and Methods). Relative fluorescent intensities of granules of a cell were calculated by dividing the average fluorescence intensity of each granule minus the background by the average fluorescence intensity of the cytoplasm minus the background. The Steel-Dwass test, a nonparametric multiple comparison test, was used to examine significant differences among the groups. *p<0.05, **p<0.005, ***p<0.0005. (D) Analysis of fluorescent intensities of Caprin1-containing granules. Fluorescent intensities of Caprin1-containing granules, the cytoplasm, and background in (B) were measured. Relative fluorescent intensities of Caprin1-containing granules were calculated in the same manner as in (C). The Steel-Dwass test was used to examine significant differences among the groups. *p<0.05, **p<0.005, ***p<0.0005, n.s.: not significant

To elucidate the contributions of individual FMRP domains/regions to RNA granule formation, we utilized several FMRP mutants carrying mutations in distinct domains/regions. Among the FMRP mutants examined, three carry FXS-associated mutations (R138Q in KH0, G266E in KH1, and I304N in KH2) (Suhl and Warren, 2015; Prieto et al., 2019). These mutations have been reported to impair distinct functions of FMRP (Feng et al., 1997; Ascano et al., 2012; Myrick et al., 2014, 2015). In another mutant, 4R-4F, four arginine residues within the RGG box that are critical for binding to G-quadruplex RNA (Blackwell et al., 2010; Prieto et al., 2019) were replaced by four phenylalanine residues. Using these mutants, we examined how mutations in these domains/regions affect the accumulation of FMRP in stress granules and neuronal transport granules. Our results showed that the G266E mutant reduced accumulation of FMRP within stress granules in HEK293T cells which express endogenous FMRP, and that both the G266E and I304N mutations suppressed accumulation of FMRP within neuronal transport granules in FMRP-deficient neurons. These findings suggest that mRNA binding mediated by the KH1 and KH2 domains is important for FMRP accumulation in RNA granules.

## Materials and Methods

### FMRP mutants and plasmids

Based on mutations found in FXS patients, we constructed FMRP variants with the R138Q, G266E, and I304N mutations; we also constructed the FMRP 4R-4F (R533F/R538F/R543F/R545F) variant, in which four arginine residues in the RGG box were replaced with four phenylalanine residues. To construct these variants, we used a pcDNA3.1 vector (pcDNA3.1-3x Flag-FMRP) containing the coding sequence of mouse FMRP and a 3x Flag tag as a template for inverse PCR. The following primers were used to construct FMRP mutants (The bolded bases indicate mismatched bases introduced to create mutations). R138Q: **G**CAAATGTGTGCCAAAGAATCAGCACA and **T**GTAAATCTTCTGGCACCTCCAGCTTG; G266E: **A**AGAGGATCAAGATGCAGTGAAAAAGGCTAGAAG and CATAAATATGAAATGTGCAGGTATCCTCATCTAAATCAATAGCAGTG; I304N: **A**TCAAGAGATCGTGGACAAGTCAGGAGTTG and TCAGCTTTCCATTTTTTCCTATTACTTTGCCTACTAAGTTTC. In the case of 4R-4F, R533F/R538F was at first constructed using the following primers: TGGAGGAGGAGGA**TTT**GGACAAGGAGGAAGAGGAAGAGGAGGAGGCTTC and **AA**CCGCCGTCCGTCTCCTCTGCGCAG. Next, the R533F/R538F mutant was the template to produce two additional mutations (R543F/R545F) using the following primers: GA**TTT**GGAGGAGGCTTCAAAGGAAACGACGATCATTCC and C**GAA**TCCTCCTTGTCCAAATCCTCCTCCTCCAC.

### Western blot

HEK293T cells were transfected with the plasmid expressing 3x Flag-tagged mouse FMRP wild type (Flag-FMRP WT) or with one of these 4 mutants using polyethyleneimine MAX (MW 40,000, Polyscience Inc). After 2 days, the cells were harvested using cell scrapers in lysis buffer (50 mM Tris-HCl (pH 7.4), 150 mM NaCl, 1% Triton X-100, 1 mM EDTA, and cOmplete protease inhibitor cocktail (Merck)). Proteins in cell lysates were separated by 10% SDS-PAGE, and fractionated proteins were transferred to a Polyvinylidene Difluoride (PVDF) membrane (Millipore). The mouse anti-Flag antibody (M2, Sigma) was used as a primary antibody at 1/2000. The membrane was washed and incubated with peroxidase-conjugated goat anti-mouse IgG (Rockland) as a secondary antibody. The chemiluminescence of the membrane was detected using a LAS-4000 image analyzer (Fujifilm). To normalize expression levels of FMRP WT and mutants relative to β-actin, the PVDF membrane was stripped and re-probed using anti-β-actin antibody (AC-74, Sigma). The chemiluminescence of the β-actin signal was measured by the image analyzer as described above. Relative expression levels of 3x Flag-FMRP variants were calculated based on the ratio of the fluorescent intensity of each FMRP variant band to that of the β-actin band.

### Induction of stress granules in HEK293T cells

HEK293T cells were transfected with the plasmid expressing Flag-FMRP WT or one of these 4 mutants using ViaFect transfection reagent (Promega). One day later, the transfected cells were harvested by trypsinization and centrifugation, and the cells were seeded on coverslips at a density of 1.8 x 10^4^ cells/cm^2^ and cultured for an additional 2 days. The transfected cells were subjected to oxidative stress treatment (200 μM sodium arsenite (As)) for 1 h, then fixed with 4% paraformaldehyde (PFA) in phosphate-buffered saline (PBS).

### *Fmr1*-KO mice and cortical culture

We used WT and *Fmr1*-KO (knockout) mice on an ICR background, which were established by backcrossing C57BL/6J *Fmr1*-KO mice (Parvin et al., 2019). All procedures were performed according to the guidelines outlined in the Institutional Animal Care and Use Committee of Yokohama City University. For mouse cortical culture, male E16 embryos of WT (*Fmr1* (+/y)) and KO (*Fmr1* (-/y)) littermates were obtained by crossing WT males (*Fmr1* (+/y)) and the heterozygous females (*Fmr1* (+/-)). Mouse cortices from WT and KO embryos were dissected and trypsinized. The resulting cells were seeded on poly-L-lysine-coated coverslips at a density of 3.2 x 10^4^ cells/cm^2^ and cultured in a CO_2_ incubator. The cortical culture was transfected with a pcDNA3.1-based plasmid harbouring the coding region of FMRP WT or one of the above mutants and EGFP connected by IRES (internal ribosome entry site) sequence using ViaFect transfection reagent (Promega), three days after plating. These plasmids were able to co-express one of the FMRP variants and EGFP simultaneously. The transfected cells were cultured for an additional 3 days and fixed with 4% PFA in PBS.

### Immunofluorescence

Immunofluorescence of fixed cells (HEK293T cells and cortical neurons) was performed as described previously. Briefly, the fixed cells were permeabilized with Tris-Buffered Saline (TBS; 50 mM Tris-HCl (pH 7.4) and 150 mM NaCl) containing 0.3% Triton X-100 for 5 min. After 1 h blocking with immunofluorescence (IF) buffer (0.1% Triton X-100 and 2% bovine serum albumin (BSA) in TBS), cells were then incubated for 1 h with primary antibodies: mouse anti-Flag antibody (1/2,000, M2, Sigma), rabbit anti-Caprin1 antibody (1/2,000, Proteintech), chick anti-GFP antibody (1/2,000, Aves Labs), rabbit anti-G3BP1 antibody (1/1,000, Sigma). After washing four times with IF buffer, the fixed cells were incubated for 30 min with fluorophore-conjugated secondary antibodies: CF488A-goat anti-chicken IgY (1/1000, Biotium), Alexa Fluor 555-Donkey Anti-mouse IgG (Jackson ImmunoResearch), Alexa Fluor 647-Donkey Anti-rabbit IgG (Jackson ImmunoResearch). Nuclei of neurons were stained with DAPI (4’,6-diamidino-2-phenylindole, 10 μg/mL, Fujifilm Wako).

### Microscopy and analysis of images

Images of IF were captured using an inverted microscope (Nikon Eclipse Ti-E) with an iXON3 CCD camera (Andor Technology) using a 60x oil-immersion lens. Analysis of IF images was performed using ImageJ/FIJI (NIH). In the case of stress granules of HEK293T cells stained with anti-Flag and Caprin1 antibodies, the interior of the cell and the exterior of the nucleus were outlined by the Polygon Selection tool of ImageJ to define cytosolic regions at first. Next, to remove fluorescence in the cytoplasm not originating from stress granules, a subtraction process was performed using the “Subtract Background” tool (rolling size = 3.2 pixels). Granules were defined using the “Analyze Particles” tool (size (micron²) = 0.10 to infinity) on images that had been binarized using the “Threshold” tool with the “yen” algorithm. Fluorescent intensities of each defined granule in a cell were measured, and the relative fluorescent intensity of granules of a cell was calculated by dividing the average fluorescence intensity of each granule minus the background by the average fluorescence intensity of the cytoplasm (excluding the nucleus and granules) minus the background.

In the case of neuronal transport granules of cortical neurons stained with anti-Flag antibody, using the “Segmented Line” tool, a line was drawn along the longest neurite from a point 20 µm away from the cell body to a point 40 µm further along, and the line was subsequently straightened using the “Straighten” tool. Next, to remove fluorescence in the cytoplasm not originating from neuronal transport granules, a subtraction process was performed using the “Subtract Background” tool (rolling size = 5.0 pixels). Neuronal transport granules were defined using the “Analyze Particles” tool (size (micron²) = 0.10 to infinity) on images that had been binarized using the “Threshold” tool with the “yen” algorithm. Fluorescent intensities of each defined granule within a neurite were measured, and relative fluorescent intensity of granules of a neurite was calculated, in the same manner as for stress granules.

## Results

### The KH1 domain mutation (G266E) reduced accumulation of FMRP within stress granules under oxidative stress

To investigate the roles of domains/regions of FMRP in RNA granule formation, we constructed four FMRP mutants (Figure 1A). Among the four mutants, three (R138Q, G266E and I304N) are based on the mutations found in FXS patients. The G266E and I304N mutants exhibit reduced binding to mRNA (Ascano et al., 2012; Myrick et al., 2014), while R138Q retains mRNA binding (Myrick et al., 2015). The 4R-4F mutant was generated by replacing four arginine residues within the RGG box with phenylalanine residues; these arginine residues have been shown to be important for binding to G-quadruplex RNA (Blackwell et al., 2010).

To examine the effect of the mutations on FMRP-containing stress granules, HEK293T cells were transfected with the plasmid expressing 3x Flag-tagged FMRP wild type (Flag-FMRP WT) or one of the Flag-FMRP mutants (R138Q, G266E, I304N, or 4R-4F). The transfected cells were subjected to oxidative stress treatment using 200 μM Sodium Arsenite (As) for 1 h. In the untreated control, Flag-FMRP WT localized diffusely throughout the cytosol. Oxidative stress treatment with As induced granule formation of Flag-FMRP WT (Figure 1B), and Flag-FMRP WT was scarcely localized in the cytoplasm outside of the granules. Most of the granules containing Flag-FMRP WT were co-localized with granules containing Caprin1, an RBP associated with stress granules, indicating that Flag-FMRP WT was contained in stress granules (Figure 1B). The R138Q, I304N, and 4R-4F mutants also formed granules in the same manner as WT. The G266E mutant also formed granules; however, a significant amount of the G266E mutant was present in the cytoplasm outside of the granules, unlike the WT and other mutants. To determine the level of accumulation of FMRP in the granules quantitatively, we measured the fluorescent intensity of the granules, following image processing to semi-automatically identify the boundaries of the granules using ImageJ. Relative fluorescent intensities of granules of each cell were calculated by dividing the average fluorescence intensity of each granule by the average fluorescence intensity of the cytoplasm excluding the granules. The relative fluorescent intensities become “1” if the intensity within the granules is the same as that in the cytoplasm outside the granules. The relative fluorescence intensity of WT after As treatment (median: 8.1) was significantly higher than that of no treatment (median: 1.7), indicating accumulation of Flag-FMRP in granules (Figure 1C). However, while the relative fluorescence intensity of the G266E mutant (median: 2.9) after As treatment was significantly lower than that of WT, the relative fluorescence intensities of the other mutants were comparable to those of WT. These results indicate that the G266E mutation in the KH1 domain suppresses accumulation of FMRP in stress granules. Next, to examine the effect of the FMRP mutations on granule formation of Caprin1, one of the stress granule markers, we analyzed relative fluorescent intensities of Caprin1 in HEK293T cells expressing FMRP WT or one of the mutants. Under As treatment, relative fluorescent intensities of Caprin1-containing granules significantly increased compared to no treatment in HEK293T cells expressing FMRP WT or any mutants (Figure 1B). The relative fluorescent intensities in cells expressing any mutants were not significantly different from those in the cells expressing FMRP WT under As treatment (Figure 1D). These results suggest that the G266E mutation in the KH1 domain of FMRP does not affect stress granule formation per se, but rather inhibits the accumulation of the G266E mutant within stress granules.

Expression levels of FMRP WT and the mutants in HEK293T cells were examined to confirm similar expression levels among them. HEK293T cells expressing 3x Flag-tagged FMRP WT or one of the mutants were analyzed using Western blotting. No difference in the mobility of the bands recognized by anti-Flag antibody was observed among the FMRP variants (Figure 2A). Although expression levels of G266E and I304N normalized by β-actin expression (Figure 2B) were slightly higher than those of WT, no statistically significant differences were observed among expression levels of FMRP WT and the mutants (Figure 2C). These results indicate that the reduced accumulation of the G266E mutant in stress granules is not due to a decrease in the expression level of the G266E mutant.

**Figure 2.**
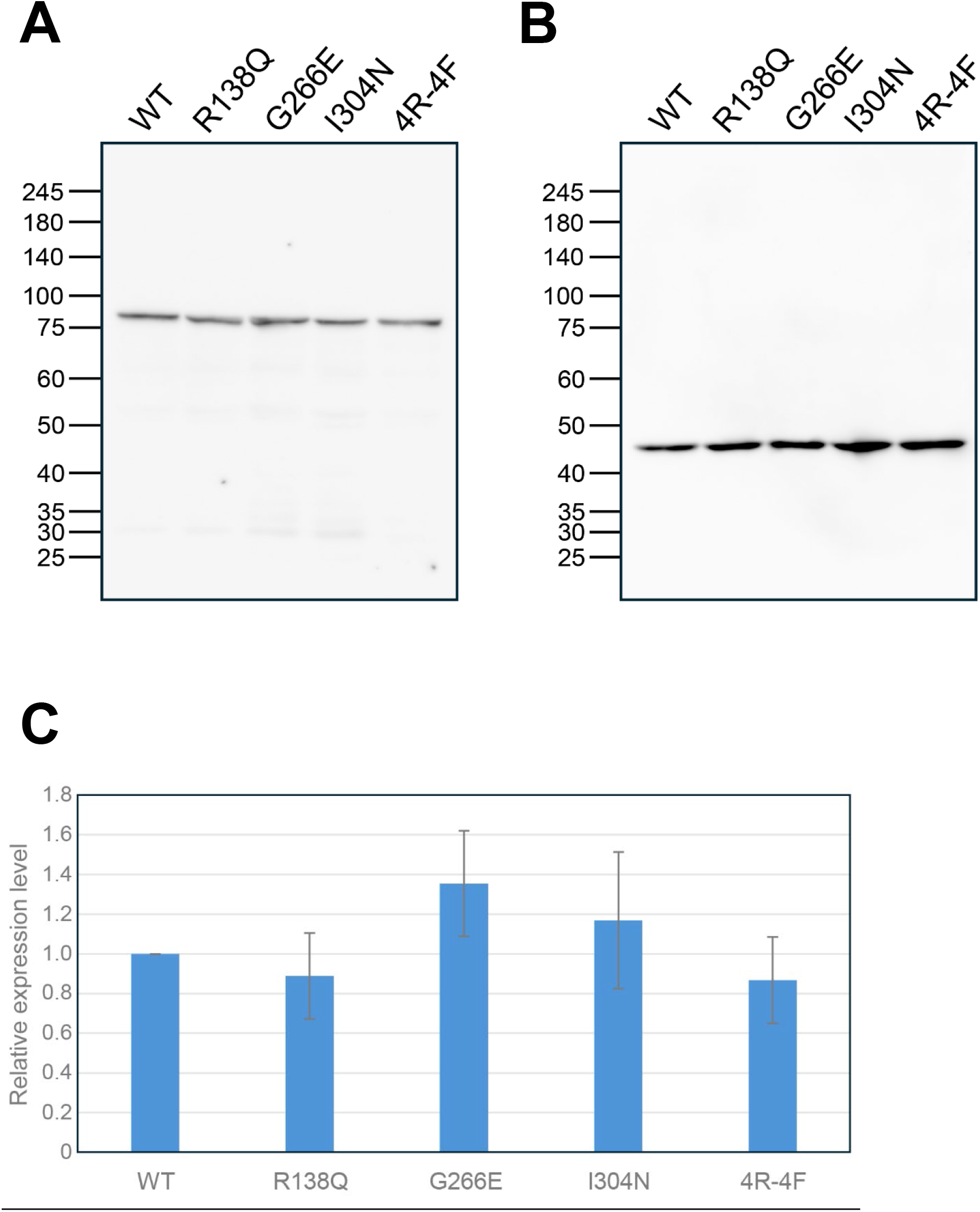
Comparison of expression levels among FMRP WT and the mutants. (A) HEK293T cells transfected with the plasmid expressing FMRP WT or one of the FMRP mutants were subjected to Western blot analysis using an anti-Flag antibody. (B) After removing the anti-Flag antibody, the membrane was re-probed with an anti-β-actin antibody. (C) Relative expression levels of 3x Flag-FMRP variants were calculated based on the ratio of the fluorescent intensity of each FMRP variant band to that of the β-actin band. Data are shown as mean ± SEM (n = 5). No statistically significant differences were detected among the groups by the Friedman test (p = 0.142).

### The KH1 (G266E) and KH2 (I304N) mutations reduced FMRP accumulation within neuronal transport granules in WT neurites

Our findings suggest that the KH1 domain is important for accumulation of FMRP within stress granules under oxidative stress (Figure 1). Next, we examined which domains/regions play an important role in accumulation of FMRP within neuronal transport granules of mouse WT cortical neurons. Cortical neurons transfected with one of the plasmids to co-express EGFP and one of the Flag-FMRP variants were fixed at DIV7 and subjected to immunofluorescence staining (Figure 3A). EGFP was used to identify cells expressing FMRP variants and visualize the shape of neurites. In neurons expressing FMRP WT, FMRP was diffusely localized in cell bodies, but within neurites, distinct granules containing FMRP were present as neuronal transport granules. FMRP-containing granules were also observed in neurites expressing the R138Q or 4R-4F mutant, similar to WT. However, in the neurite expressing the G266E mutant, localization of the FMRP mutant was more diffuse compared to the WT. Although distinct granules containing FMRP I304N were observed in a small number of neurites, the localization of the I304N mutant in most neurites was diffuse, similar to that of the G266E mutant. To calculate the relative fluorescent intensities of granules containing FMRP in neurites, the longest neurites of FMRP WT and mutants were straightened, and the fluorescent intensities of FMRP-containing granules were measured (Figure 3B). The relative fluorescent intensities of granules of each neurite were calculated by dividing the average fluorescence intensity of each granule by the average fluorescence intensity of the neurite excluding the granules. The relative fluorescence intensities of G266E and I304N were significantly lower than that of WT (Figure 3C). The relative fluorescence intensity of G266E was slightly lower than that of I304N (p = 0.0508). These results indicate that the accumulation of the G266E mutant within neuronal transport granules was markedly reduced, whereas the I304N mutant occasionally showed distinct granular localization. When the localization of FMRP WT and G3BP1, which is another RBP that serves as a component of both stress granules and neuronal transport granules, was compared in neurites, some granules contained both FMRP and G3BP1, but a significant number of granules contained only FMRP or G3BP1 (Figure 3D). The limited colocalization of FMRP and G3BP1 in neuronal transport granules contrasted with the colocalization of FMRP and Caprin1 observed in stress granules. FMRP G266E and I304N were localized diffusely and rarely co-localized with G3BP1-containing granules, while G3BP1 formed distinct granules. These results suggest that KH1 and KH2 domains are important for the accumulation of FMRP within neuronal transport granules, without affecting G3BP1 granule formation.

**Figure 3.**
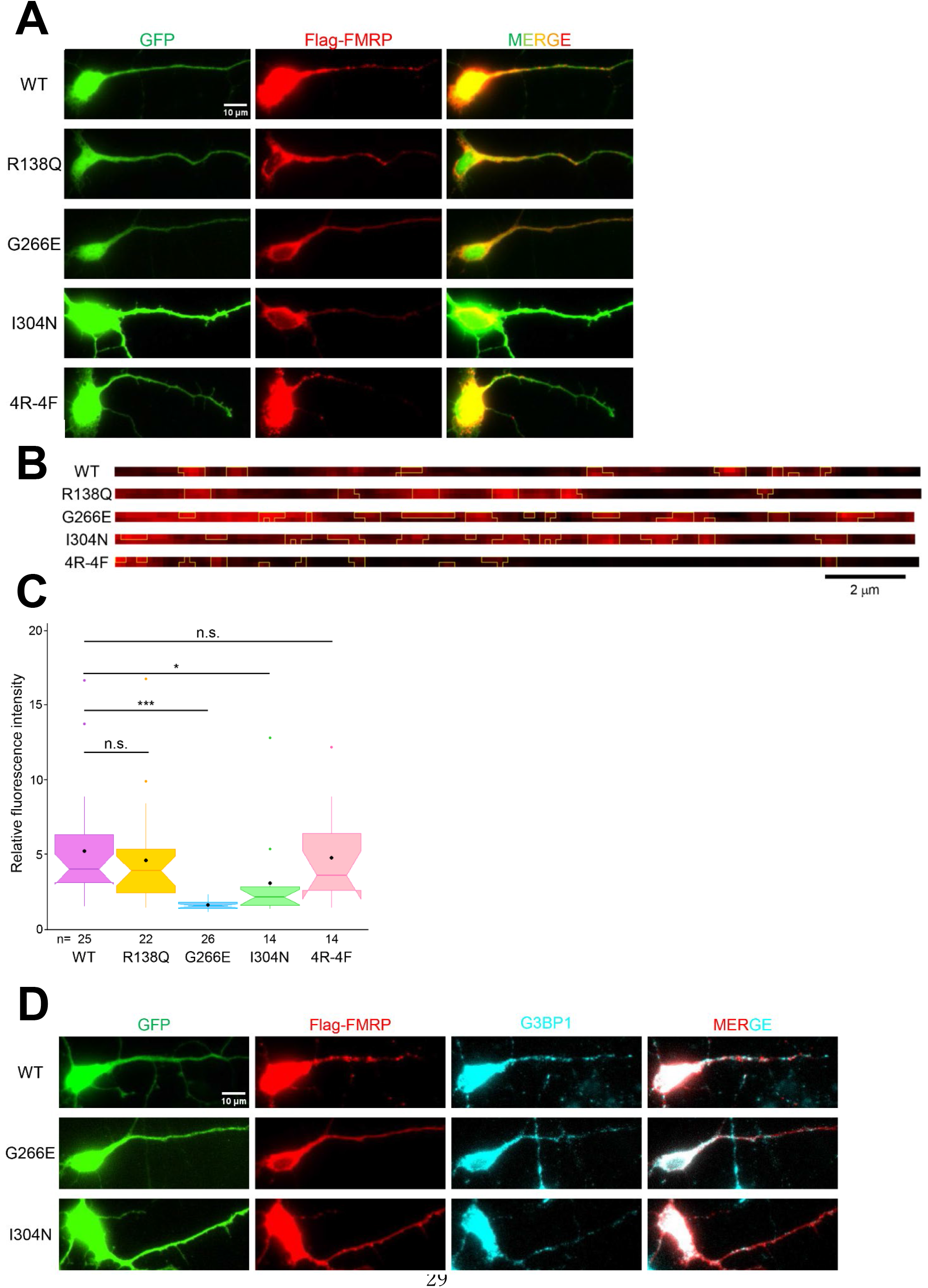
The KH1 and KH2 mutations (G266E and I304N) reduced accumulation of FMRP within neuronal transport granules in mouse cortical WT neurons. (A) Mouse cortical neurons derived from male E16 WT (*Fmr1* (+/y)) embryos were transfected with plasmids co-expressing EGFP and FMRP WT or one of the FMRP mutants (R138Q, G266E, I304N, and 4R-4F). The transfected neurons were fixed at DIV7, followed by immunostaining using anti-GFP and anti-Flag antibodies. Scale bar: 10 μm. (B) The longest neurites of the transfected neurons shown in (A) were straightened to calculate relative fluorescent intensities. A segment of the longest neurite, extending from a point 20 µm away from the cell body to a point 40 µm further along, was traced and subsequently straightened. FMRP-containing granules (pixels enclosed by the yellow line) were defined in the same manner as in Figure 1C. (C) Analysis of fluorescent intensities of FMRP-containing granules. Fluorescent intensities of FMRP-containing granules, the neurites, and the background in (B) were measured. Relative fluorescent intensities of granules of a neurite were calculated by dividing the average fluorescence intensity of each granule minus the background by the average fluorescence intensity of the neurite minus the background. The Steel-Dwass test, a nonparametric multiple comparison test, was used to examine significant differences among the groups. *p<0.05, **p<0.005, ***p<0.0005, n.s.: not significant. (D) Localization of Flag-FMRP and G3BP1 in neurites of WT neurons. Cortical neurons were transfected with plasmids co-expressing EGFP and FMRP WT or one of the FMRP mutants (G266E and I304N). Scale bar: 10 μm.

### The KH1 (G266E) and KH2 (I304N) mutations reduced FMRP accumulation within neuronal transport granules in *Fmr1*-KO neurites to a similar extent

Because endogenous FMRP is expressed in WT neurons, it is possible that localization of heterodimers formed between endogenous FMRP and the mutant FMRP expressed by transfection is different from that of homodimers of the mutant FMRP. To rule out the influence of endogenous FMRP, we used *Fmr1*-KO (knockout) neurons to express FMRP WT or one of the mutants. In KO neurons expressing FMRP WT, distinct granules containing FMRP were localized within neurites (Figure 4A). While distinct granules containing FMRP were also observed in neurites expressing the R138Q or 4R-4F mutant, localizations of the G266E and I304N mutants in *Fmr1*-KO neurites were more diffuse compared to the WT. The fluorescent intensities of FMRP-containing granules were measured in the longest neurites of FMRP WT and mutants (Figure 4B). The relative fluorescence intensities of G266E and I304N were significantly lower than that of WT (Figure 4C). The relative fluorescence intensity of G266E was almost the same as that of I304N in *Fmr1*-KO neurites, unlike that of WT neurites. The number of granules defined by ImageJ was not significantly different among WT, G266E, and I304N-expressing KO neurites (Figure 4D). Total areas of granules in G266E- and I304N-expressing KO neurites were significantly increased compared to WT (Figure 4E), indicating that FMRP-positive regions were more spatially diffuse in neurons expressing these mutants than in those expressing WT FMRP. These data suggest that both the G266E and I304N mutations reduce FMRP accumulation within neuronal transport granules without decreasing the number of FMRP-positive regions detected by image analysis. In KO neurons as well, FMRP WT partially co-localized with G3BP1, whereas both the G266E and I304N mutants rarely co-localized with G3BP1-containing granules (Figure 4F), similar to WT neurons. Our results suggest that the KH1 and KH2 domains, which are responsible for binding mRNAs, play a crucial role in the accumulation of FMRP within neuronal transport granules.

**Figure 4.**
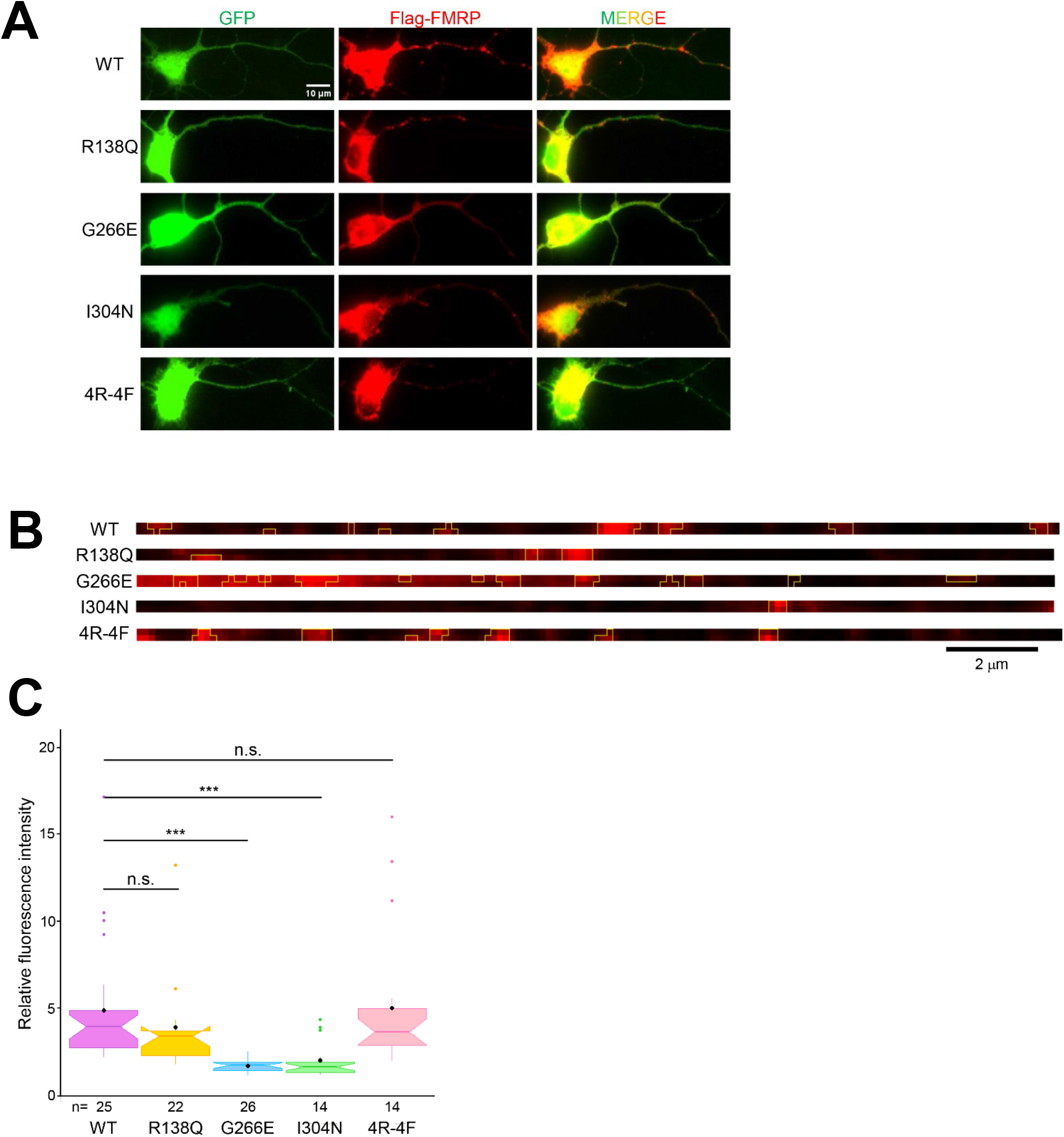
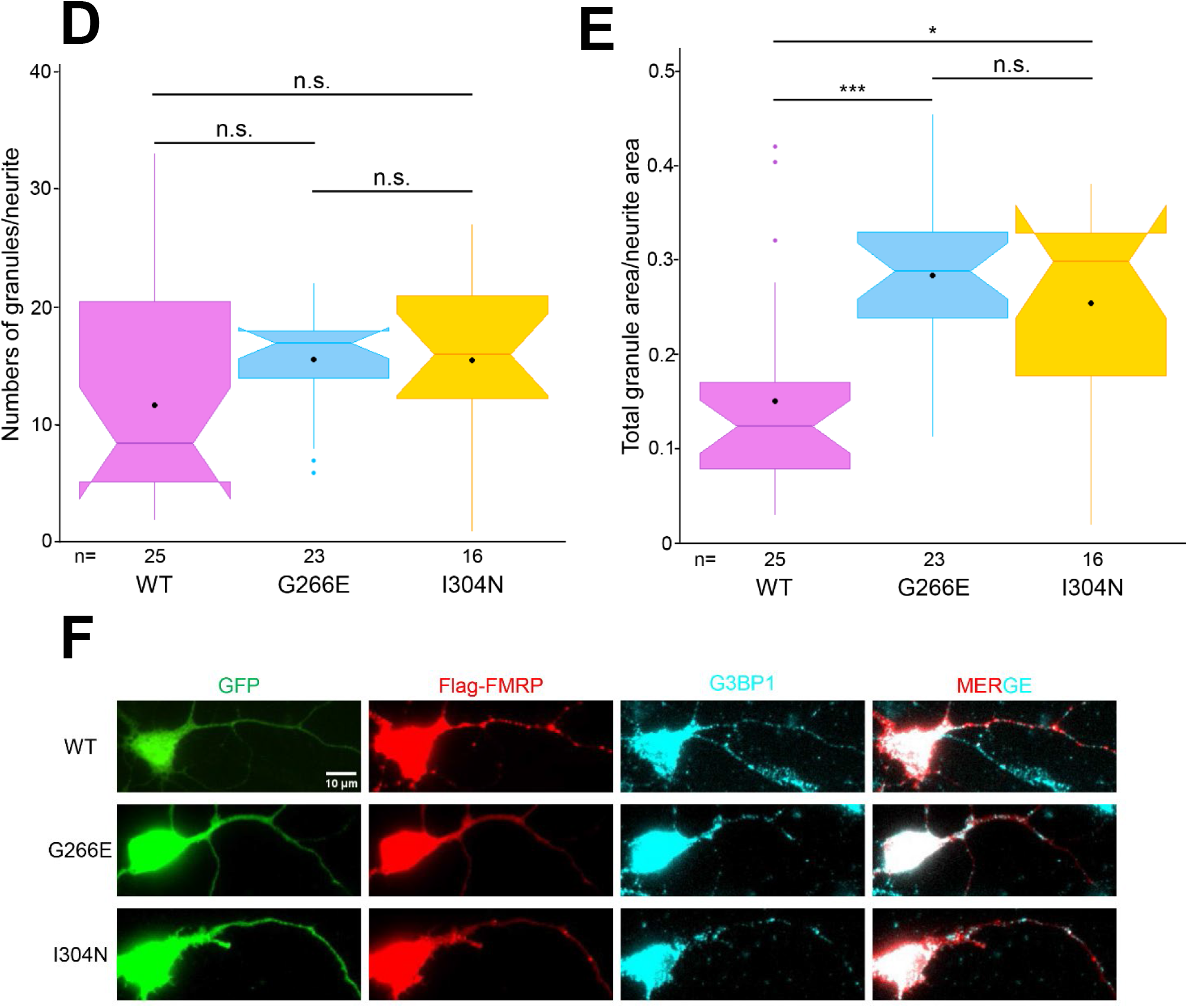
The KH1 and KH2 mutations (G266E and I304N) reduced accumulation of FMRP within neuronal transport granules in mouse *Fmr1*-KO cortical neurons. (A) Mouse cortical neurons derived from male E16 *Fmr1*-KO embryos (*Fmr1* (-/y)) were transfected with plasmids co-expressing EGFP and FMRP WT or one of the FMRP mutants (R138Q, G266E, I304N, and 4R-4F). The transfected neurons were fixed at DIV7, followed by immunostaining using anti-GFP and anti-Flag antibodies. Scale bar: 10 μm. (B) The longest neurites of the transfected neurons shown in (A) were straightened to calculate relative fluorescent intensities. FMRP-containing granules (pixels enclosed by the yellow line) were defined in the same manner as in Figure 1C. (C) Analysis of fluorescent intensities of FMRP-containing granules in *Fmr1*-KO neurons. Fluorescent intensities of FMRP-containing granules, the neurites, and background in (B) were measured. Relative fluorescent intensities of granules of a neurite were calculated in the same manner as described for Figure 3C. The Steel-Dwass test, a nonparametric multiple comparison test, was used to examine significant differences among the groups. ***p<0.0005, n.s.: not significant. (D) Number of FMRP-containing granules in *Fmr1*-KO neurites co-expressing EGFP and FMRP WT or one of the FMRP mutants (G266E and I304N). FMRP-containing granules in Figure 4B were counted, and the number of granules per neurite (40 µm long) was plotted. The Steel-Dwass test was used to examine significant differences among the groups. n.s.: not significant. (E) Total granule areas relative to neurite area in *Fmr1*-KO neurites. Total areas of FMRP-containing granules in Figure 4B per neurite were calculated and plotted. The Steel-Dwass test was used to examine significant differences among the groups. *p<0.05, **p<0.005, ***p<0.0005, n.s.: not significant. (F) Localization of Flag-FMRP and G3BP1 in neurites of *Fmr1*-KO neurons. *Fmr1*-KO neurons were transfected with plasmids co-expressing EGFP and FMRP WT or one of the FMRP mutants (G266E and I304N). Scale bar: 10 μm.

## Discussion

In this study, as a first step toward elucidating the mechanism underlying the formation of neuronal transport granules, we focused on FMRP, a component of both stress granules and neuronal transport granules. To identify FMRP domains or regions important for its accumulation within RNA granules, we examined four FMRP mutants affecting distinct domains or regions. The G266E mutation in the KH1 domain of FMRP reduced accumulation of FMRP in stress granules in HEK293T cells without affecting the formation of Caprin1-containing stress granules. In *Fmr1*-KO cortical neurons, both the G266E and I304N mutations in the KH1 and KH2 domains suppressed accumulation of FMRP within neuronal transport granules without affecting G3BP1-containing granules. These results suggest that the KH1 and KH2 domains, which possess RNA-binding activity, play important roles in the accumulation of FMRP within RNA granules.

Among the four types of FMRP mutants, only the G266E mutant showed reduced accumulation in stress granules (Figure 1). In contrast, both the G266E and I304N mutations suppressed accumulation of FMRP in neuronal transport granules in *Fmr1*-KO neurons, and the degree of suppression was comparable between the two mutants (Figure 4). In both FMRP-deficient mouse fibroblasts and FXS patient-derived lymphoblasts, the I304N mutant failed to accumulate in stress granules (Didiot et al., 2009), suggesting that accumulation of the I304N mutant in stress granules is suppressed in cells lacking endogenous FMRP. Furthermore, in WT neurons expressing endogenous FMRP, unlike *Fmr1*-KO neurons, the I304N mutant accumulated more in neuronal transport granules than the G266E mutant (Figure 3). Therefore, it is suggested that both the G266E and I304N mutants have reduced ability to accumulate within RNA granules, such as stress granules and neuronal transport granules, in the absence of endogenous FMRP. Because both the G266E and I304N mutants have decreased mRNA-binding activity (Ascano et al., 2012; Myrick et al., 2014), mRNA binding mediated by KH1 and KH2 may contribute to multivalent interactions underlying RNA granule assembly, potentially including phase-separation processes. The KH1 domain carrying the G266E mutation exhibited an unfolded structure even at 25°C (Catalano et al., 2024), while stability of the I304N mutant decreased at temperatures above 40°C (Valverde et al., 2007). These findings suggest that the G266E mutant may be structurally less stable than the I304N mutant. It is possible that an unstable domain may weaken not only multivalent interactions between FMRP and mRNAs, but also multivalent interactions with other RBPs or endogenous FMRP. The difference in stability between the G266E and I304N mutants may explain why the G266E, but not I304N, mutant reduced accumulation in RNA granules in the presence of endogenous FMRP.

Because FMRP mutants with reduced mRNA-binding activity showed impaired accumulation in both stress granules and neuronal transport granules, these two types of RNA granules may share mechanisms that mediate FMRP recruitment. On the other hand, FMRP WT-containing granules showed almost complete colocalization with stress granules containing Caprin1 (and G3BP1; data not shown) (Figure 1), although FMRP WT-containing granules and G3BP1-containing granules within neurites exhibited only partial colocalization (Figure 3). These data suggest that stress granules and neuronal transport granules share similar mechanisms of granule formation while also possessing distinct characteristics. In the process of stress granule formation, G3BP1 and G3BP2 represent central nodes in the stress granule network, and Caprin1 binding to G3BP facilitates LLPS required for the formation of stress granules (Ivanov et al., 2019; Hofmann et al., 2021). FMRP may be recruited into G3BP-driven stress granules either as an individual FMRP-mRNA complex or as preassembled FMRP-containing granules. On the other hand, it is possible that G3BP1 and FMRP may independently assemble into distinct neuronal transport granules in neurites, and that subsequent interactions or co-assembly between these populations may generate granules containing both FMRP and G3BP1. The G266E mutant may have a reduced ability to accumulate within or interact with other RNA granule assemblies, while the I304N may still possess the ability to incorporate other granules via endogenous FMRP and other RBPs. In *Drosophila* motor neurons lacking FMRP, the G269E mutant (corresponding to the mouse G266E mutant) showed no localization within neurites, whereas the I307N mutant (corresponding to the mouse I304N mutant) formed distinct RNA granules within neurites, albeit in small numbers (Starke et al., 2022). These results were different from our findings in mice, where both the G266E and I304N mutants were diffusely localized within the neurites of *Fmr1*-KO cortical neurons (Figure 4). This discrepancy may be due to differences in species or cell types.

FXS-associated FMRP mutations impair the ability of FMRP to assemble into RNA granules, although the extent of impairment differs according to the mutation and cellular context. Importantly, *de novo* variants in multiple genes involved in stress granule assembly, including *G3BP1*, *G3BP2*, *UBAP2L*, and *CAPRIN1*, have been associated with neurodevelopmental disorders, and several of these variants disrupt stress granule formation (Jia et al., 2022). Our findings extend this concept to FMRP and suggest that defective RNA granule assembly may represent a shared molecular mechanism underlying genetically distinct neurodevelopmental disorders. Together with previous evidence linking mutations in stress granule-associated genes to neurodevelopmental disorders, our findings raise the possibility that disruption of RBP assembly into RNA granules, including neuronal transport granules, may represent a convergent molecular mechanism contributing to neurodevelopmental disorders.

## Acknowledgements

This work is partly supported by JSPS Grant-in-Aid for Scientific Research (KAKENHI) (C) (No. 20K06877, 23K05964) (Y. Sasaki). We thank Mr. Yoshinari Fujiwara for helpful discussions and technical assistance.

